# Time-lapse mechanical imaging of neural tube closure in live embryo using Brillouin microscopy

**DOI:** 10.1101/2022.10.06.511204

**Authors:** Chenchen Handler, Giuliano Scarcelli, Jitao Zhang

**Affiliations:** Fischell Department of Bioengineering, A. James Clark School of Engineering, University of Maryland, College Park, MD 20742, USA; Biomedical Engineering Department. Wayne State University, Detroit, MI 48201, USA

**Keywords:** neural tube closure, embryo, biomechanics, Brillouin microscopy

## Abstract

Neural tube closure (NTC) is a complex process of embryonic development involving molecular, cellular, and biomechanical mechanisms. While the genetic factors and biochemical signaling have been extensively investigated, the role of tissue biomechanics remains mostly unexplored due to the lack of tools. Here, we developed a new optical modality that can conduct time-lapse mechanical imaging of neural plate tissue as the embryo is experiencing neurulation. This technique is based on the combination of a confocal Brillouin microscope and an on-stage incubator for the modified *ex ovo* culturing of chick embryo. With this technique, for the first time, we captured the mechanical evolution of the neural plate tissue with live embryos. Specifically, we observed the continuous stiffening of the neural plate during NTC for *ex ovo* cultured embryos, which is consistent with the data of *in ovo* culture as well as previous studies. Beyond that, we found the tissue stiffening was highly correlated with the tissue thickening and bending. We foresee this non-contact and label-free technique can open new opportunities to understand the biomechanical mechanisms in development.

**Summary statement:** An all-optical technique is developed to capture the evolution of tissue mechanics during neural tube closure of live chick embryo.

## INTRODUCTION

Neural tube closure (NTC) is a central procedure of vertebrate neurulation where the planar neural plate will be elevated and fused to form a hollow neural tube. A failure of this procedure can result in severe neural tube defects, which represent one of the most common human birth defects (Wallingford et al., 2013). Genetic and molecular processes that guide NTCs have been extensively studied for many decades (Colas and Schoenwolf, 2001; Copp et al., 2003; Wilde et al., 2014). On the other hand, biomechanical mechanisms that may be involved in NTCs are attracting increasing attention in recent years (Koehl, 1990; Schoenwolf and Smith, 1990; Vijayraghavan and Davidson, 2017). On cell and tissue level, the morphogenesis of neural tube can be considered as a result of the interaction between the generated force and the mechanical resistance of the embryonic tissue (Heer and Martin, 2017; Nikolopoulou et al., 2017): the successful closure of the neural tube requires the intrinsic force can overcome the opposing tissue tension that relies on its elastic property. As such, the alteration of tissue biomechanics can cause the failure of the closure and thus malformation of neural tube (Galea et al., 2017). Although the force production and mechanical change of tissue during the procedure of NTC have been observed in experiments (Zhou et al., 2009; Zhou et al., 2015; Galea et al., 2017), the quantitative contribution of specific biomechanical processes to ensure robust neurulation remains mostly unknown. One of the main reasons is the lack of tools that can map the biomechanics of neural plate tissue *in situ* and in real time when the embryo is developing.

Many important techniques have been developed to quantify the mechanical properties of embryonic tissue (Campas, 2016), which can be approximately classified into three categories: (1) contact-based techniques, including atomic force microscopy (AFM) (Franze, 2011; Barriga et al., 2018) or microcantilever (Zhou et al., 2009; Chevalier et al., 2016; Marrese et al., 2019) based indentations, micropipette aspiration (Wen et al., 2015), and tensile test (Wiebe and Brodland, 2005). While the contact-based techniques can provide direct quantification of tissue’s elastic modulus, they need physical access of the sample and to apply force to deform the sample during measurement. Since neural tube tissue has irregular shape in 3D and mechanically interconnected, isolate explants are usually required for unambiguous mechanical test. (2) Bead/droplet-based sensors, including optical/magnetic tweezer (Welte et al., 1998; Savin et al., 2011) and microdroplet (Campàs et al., 2014). Optical/magnetic tweezer uses force-driven rigid bead to sense the mechanical properties of localized tissue, and microdroplet uses deformable droplet to quantify the tissue stress. These sensors can quantitatively measure the mechanical properties with subcellular resolution after careful calibration. However, they require injection of beads or droplets into tissue, making them invasive and low throughput. (3) Tissue ablation/dissection. This method uses either an ultrafast pulsed laser beam (Galea et al., 2017) or a blade (Beloussov et al., 1975) to dissect a portion of the tissue and evaluate the mechanical properties based on the relaxation response. This is an attractive technique because of the simple setup. However, due to the mechanical connection of embryonic tissue in 3D, this method mostly provides global assessment on relatively large scale. To summarize, existing methods have greatly advanced the assessment of embryonic tissue biomechanics. However, due to the technical limitations, the *in situ* mechanical mapping of the neural plate tissue during the procedure of NTC in live embryo has not been reported.

Confocal Brillouin microscopy is an emerging technique for quantifying the mechanical properties of biological materials (Scarcelli et al., 2015; Prevedel et al., 2019; Zhang and Scarcelli, 2021). Different from conventional mechanical test methods, Brillouin microscopy uses a laser beam to measure the elastic properties of the material. This is based on an optical phenomenon called spontaneous Brillouin light scattering (Boyd, 2003), where the interaction of the incident laser beam and the acoustic phonon within the material will introduce a frequency shift (i.e., Brillouin shift) to the scattered light (see Material and methods). By measuring the Brillouin shift of the scattered light using a customized spectrometer, the elastic longitudinal modulus of the material can be directly quantified. Since Brillouin microscope is designed in a confocal configuration, it can achieve diffraction-limited spatial resolution. In the past several years, we have innovated this technique and demonstrated its feasibility for quantifying the mechanical properties of single cell (Wisniewski et al., 2020; Zhang et al., 2020), embryonic tissue (Raghunathan et al., 2017), and neural plate (Zhang et al., 2018) with subcellular resolution and enough mechanical sensitivity. As an all-optical technique, Brillouin microscope can conduct measurement in a non-contact, non-invasive, and label-free manner. Therefore, it could be a promising tool for mapping the elastic property of neural plate tissue *in situ* during embryonic development.

In this work, we developed a new modality that can conduct time-lapse mechanical imaging of neural tube closure in live chick embryo. To do so, we integrated a confocal Brillouin microscope with an on-stage incubator for modified *ex ovo* culturing. The *ex ovo* culture ensures the embryo continuously develops over 21 hours, covering the complete events of NTC. We then used Brillouin microscope to acquire 2D mechanical images of neural plate tissue as the embryo was experiencing neurulation. With this new modality, we observed a distinct increase in the averaged Brillouin shift of the neural plate tissue during the NTC of the *ex ovo* cultured chick embryo, which is consistent with the results of the *in ovo* culturing system. Importantly, we found the tissue stiffening on dorsal-ventral axis was strongly correlated with the thickening of the neural plate as well as the closure angle, indicating the tissue mechanics may be synchronized with the geometric change of the neural plate to achieve a successful closure of neural tube. Together, this time-lapse mechanical imaging modality can provide new data for understanding the biomechanical mechanisms during embryonic neurulation.

## RESULTS

Time-lapse 2D mechanical imaging was performed on an inverted Brillouin microscope (**Fig. 1A**) (see Materials and methods). By definition, Brillouin shift is positively linked to the longitudinal modulus by material properties including refractive index *n* and density *ρ*. For biological materials, the ratio of refractive index and density *ρ/n*^2^ is found to be approximately constant (Scarcelli et al., 2013; Scarcelli et al., 2015). Therefore, we here used the Brillouin shift to interpret the relative change of elastic modulus. In the modified *ex ovo* culture, the embryo was placed into a petri dish with dorsal side facing down (**Fig. 1B**). The dish was then placed into the on-stage incubator for sustaining the development of the embryo (>21 hours). The time-lapse bright-field images suggest the embryos from *ex ovo* culture have developed with the similar time rate as those from *in ovo* culture (**Fig. S1–S2**).

**Figure 1.**
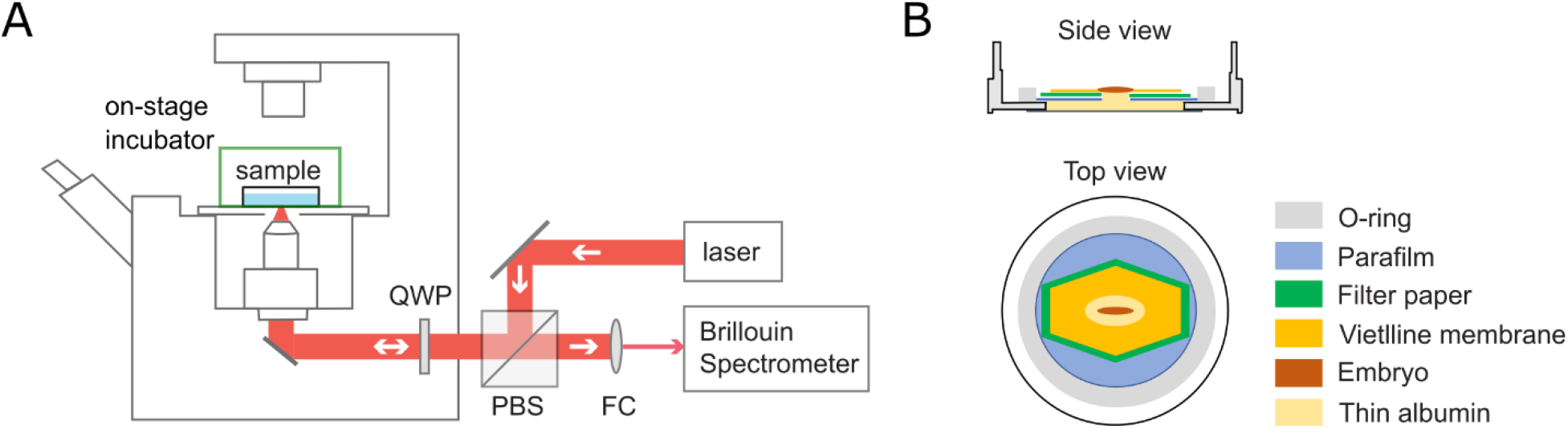
Schematic of the setup. (A) Confocal Brillouin microscope with on-stage incubator. QWP: quarter-wave plate; PBS: polarized beam splitter; FC: fiber coupler. (B) Carrier dish for *ex ovo* culture. Side view (top) and top view (bottom) displays all components including embryo within the carrier.

### *In ovo* cultured embryos show increased Brillouin shift of neural plate against developmental stage

To exclude any potential impact of the *ex ovo* culture and the laser illumination on the tissue mechanics of neural plate, we collected *in ovo* cultured embryos (N=46) at different Hamburger Hamilton (HH) stages (HH 6 to HH 12) and acquired 2D mechanical images of the cross-section perpendicular to the anterior-posterior axis using Brillouin microscope. The representative Brillouin images suggest that the neural plate of the embryo at later stage has higher Brillouin shift than that at earlier stage (**Fig. 2A-2D**). We then quantified the average Brillouin shift of the neural plate region for all the collected embryos. We observed that the Brillouin shift of the neural plate showed a distinct increase from HH 6 to HH 9+ and approximately maintained its value afterward (**Fig. 2E**). The neural plate of later-stage embryo (i.e., HH 12) has an average Brillouin shift of 6.353 GHz, which is 0.126 GHz higher than that of early-stage embryos (i.e., HH 6), corresponding to ~60% increase of Young’s modulus according to the empirical relationship between longitudinal modulus and Young’s modulus obtained from cells (Zhang et al., 2020) (see Materials and methods).

**Figure 2.**
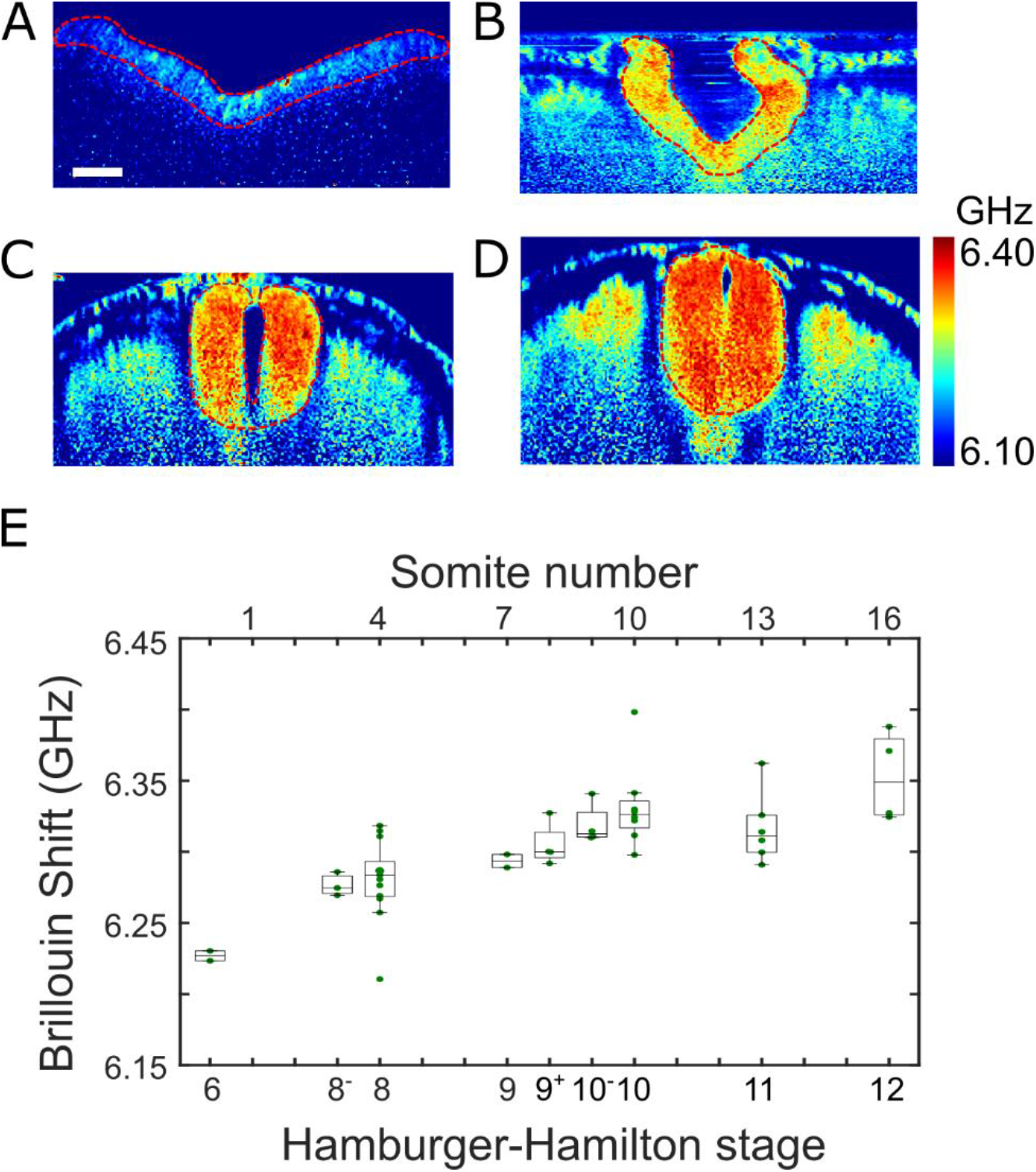
Results of *in ovo* cultured embryos. (A)-(D) Representative of four different embryos at different HH stages: (A) HH 6, (B) HH 8, (C) HH 11, (D) HH 12. Red dashed line outlines the neural plate region. (E) Average Brillouin shifts of neural plate at different stages reveal continual tissue stiffening. Number of embryos: 46. Scale bar is 50 μm.

### Time-lapse mechanical imaging of *ex ovo* cultured embryo shows stiffening and thickening of the neural plate during NTC

To quantify the mechanical evolution of the neural plate during the entire procedure of NTC, we conducted time-lapse mechanical mapping of *ex ovo* cultured embryo. The embryo was cultured for more than 14 hours (**Fig. S3**), within which the time-lapse Brillouin image of the neural plate cross-section was acquired at the hindbrain/cervical region (**Fig. 3A**). The result shows the average Brillouin shift of the neural plate continuously increases with culturing time (**Fig. 3B**), which is consistent with the result of *in ovo* cultured embryos. Repeat experiments (N=9) suggest the stiffening of the neural plate during NTC is a common phenomenon for chick embryos (**Fig. 3D**). Specifically, the Brillouin shift of the neural plate increased significantly from HH 8- to HH 9 and remained minor change afterward. At the endpoint of *ex ovo* culture (HH 10), the neural plates have an average Brillouin shift of 6.336 GHz, which is 0.097 GHz higher than that of the earliest stage (HH 8-), corresponding to the relative increase of ~ 46% in terms of the Young’s modulus. This is consistent with the result of *in ovo* cultured embryos, confirming that the *ex ovo* culture and laser illumination did not affect the mechanical evolution of the neural plate tissue during embryonic development.

**Figure 3.**
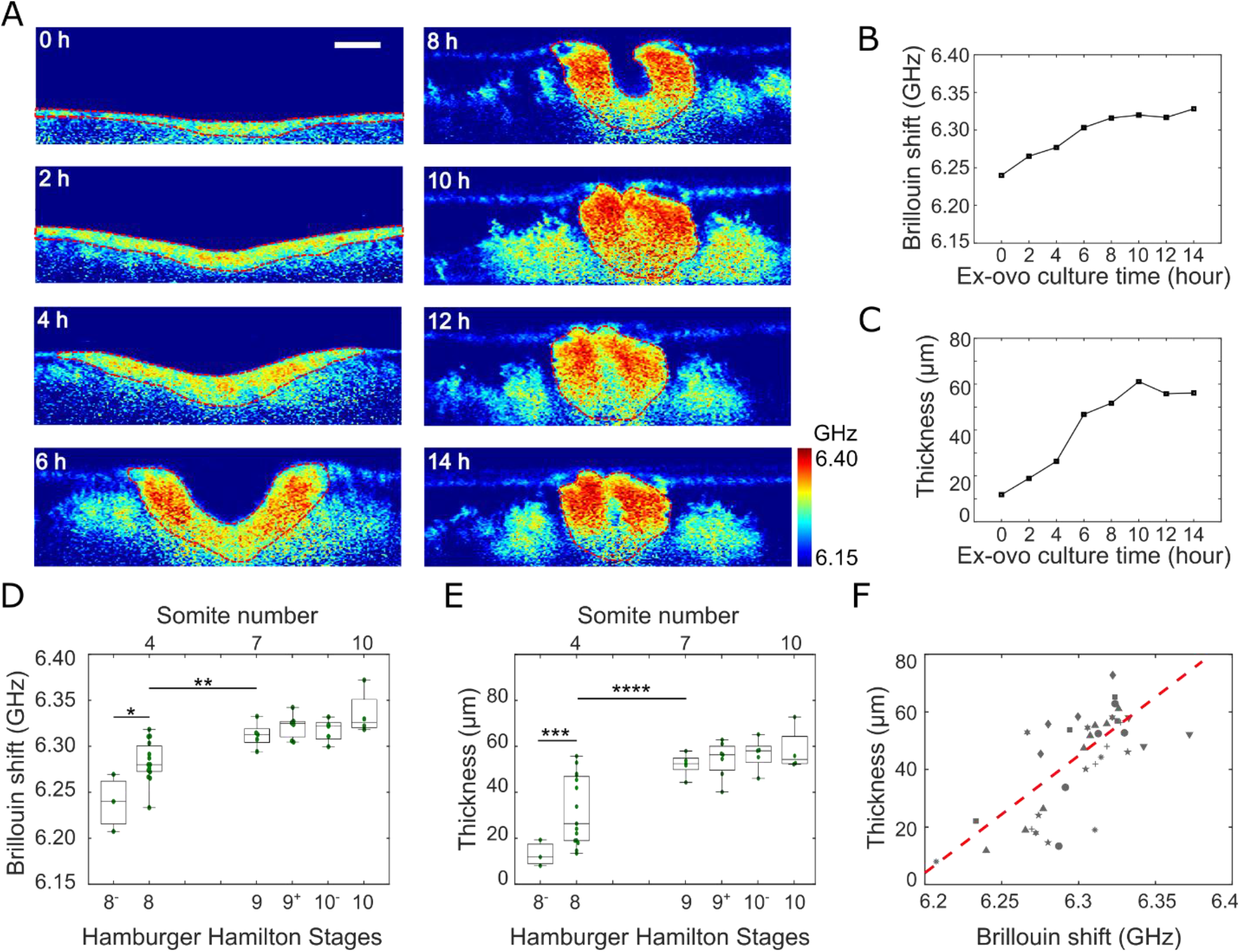
Time-lapse mechanical imaging of *ex ovo* cultured embryos. (A) Representative time-lapse Brillouin images of an embryo over 14 hours. (B) Average Brillouin shift of neural plate tissue of the embryo in (A) is increasing with culturing time. (C) Thickness of neural plate tissue of the embryo in (A) is increasing with culturing time. (D) Tissue stiffening against developmental stage is observed for all the embryos. (E) Tissue thickening against developmental stage is observed for all the embryos. (F) Correlation between average Brillouin shift and thickness of neural plate. Symbols represent different embryos. Number of embryos: 9. Number of measurements: 39. Two-sample *t* test is used to quantify the statistical significance. \**p*=0.008; ** *p*=0.012; *** *p*=0.049; **** *p*=0.01. Scale bar is 50 μm.

Using tissue mechanics as a contrast mechanism in Brillouin imaging, we can also quantify the morphological change of the neural plate during NTC. Here we measured the averaged thickness of the two sites that were in the middle of the distance between the median hinge point and the tips (**Fig. 3C**). Consistent with published literatures (Colas and Schoenwolf, 2001; Lowery and Sive, 2004), we observed 4-fold thickening of the neural plate from HH 8- (~13 μm) to HH 9 (~52 μm) (**Fig. 3E**). Intriguingly, we found that the tissue thickening and the tissue stiffening exhibit very similar trend during the procedure of NTC. We then plotted the Brillouin shift against the thickness for all the *ex ovo* cultured embryos and found a strong correlation (*p* < 1 × 10^-6^) between them (**Fig. 3F**). This data suggests that the tissue stiffening and thickening are probably coordinated events for NTC.

### Tissue stiffening is correlated with the closure angle of neural plate during NTC

NTC is a complex biomechanical process of tissue shaping and patterning that are driven by force and mechanical properties of the tissue. Therefore, it is fundamentally necessary to understand the relationship between tissue mechanics and geometry. Here, we investigated how the tissue stiffening is related to the geometric change of the neural plate. According to the Brillouin images, we defined the closure angle *β* as the intersection of the left- and right-side neural plates at the median hinge point (**Fig. 4A**). Next, we distributed *ex ovo* cultured embryos into each 10° interval and calculated the averaged values for any interval having multiple embryos. We then plotted the Brillouin shift *ω_B_* against the closure angle *β* (**Fig. 4B**). The data can be well fitted by a simple exponential curve *ω_B_* = *A* · exp(*B* · *β*) + *C*, with fitted parameters *A* = −0.024, *B* = 7.85 × 10^−3^, and *C* = 6.348. This exponential relationship suggests that the tissue stiffening is probably synchronized with the bending of the neural plate during the procedure of NTC.

**Figure 4.**
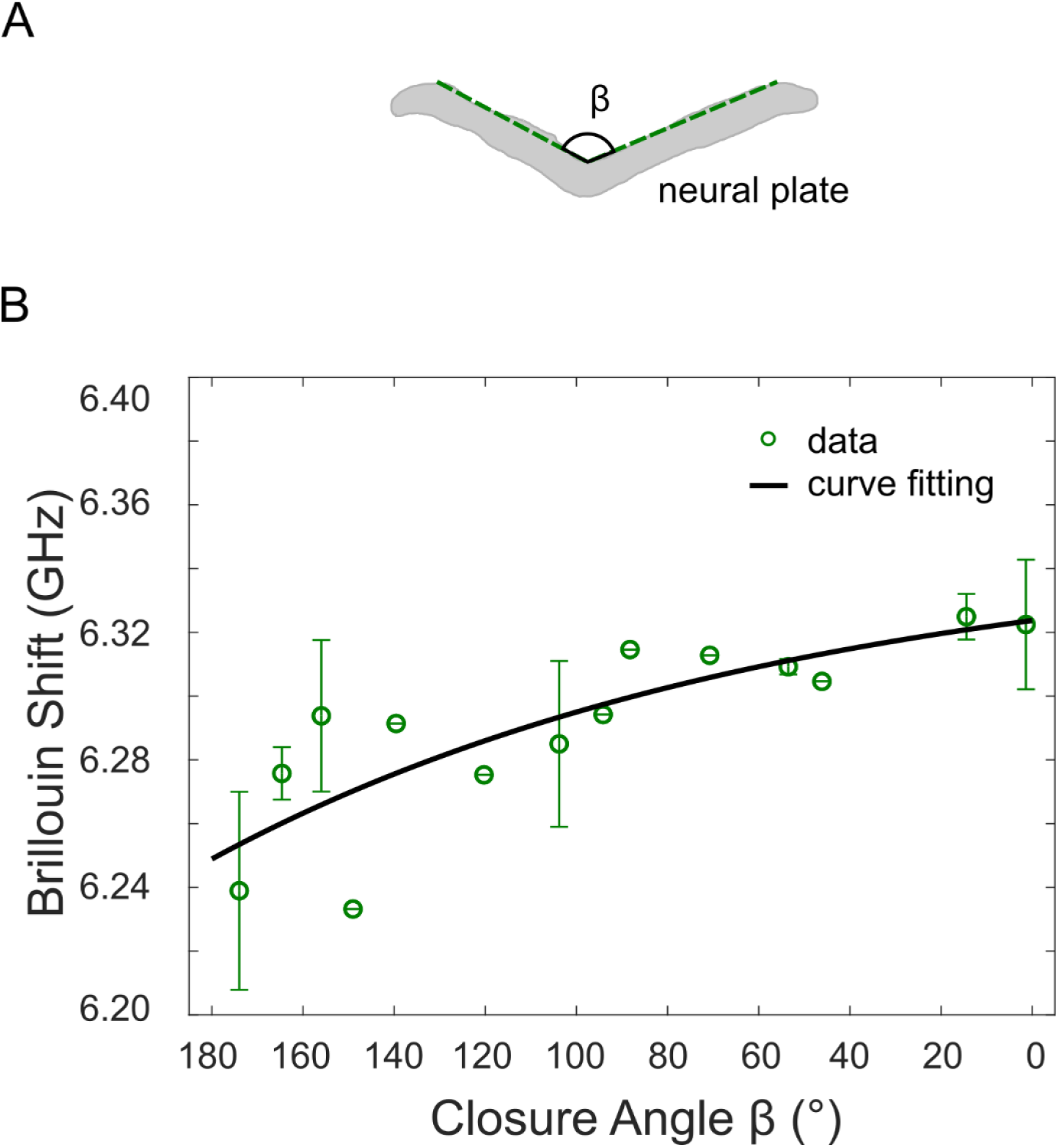
Tissue stiffening is correlated with tissue bending for *ex ovo* cultured embryos. (A) Definition of the closure angle. (B) Closure angle is correlated with average Brillouin shift of neural plate. Embryos are distributed into each 10° interval based on the closure angle. Data point represents the average value of the embryos within the same interval. Error bar represents the standard deviation. Solid curve is fitting result of an exponential function.

## DISCUSSION

Here, we developed a new imaging modality for time-lapse mechanical mapping of live chick embryo. This modality is based on the combination of a confocal Brillouin microscope and a modified *ex ovo* culturing system, which has subcellular resolution and enough mechanical sensitivity. Different from conventional techniques for mechanical test, our method used a focused laser beam to quantify the tissue mechanics, making it non-contact, non-invasive, and label free. We confirmed that the *ex ovo* culture and laser illumination did not disturb the development of the embryo. We demonstrated the feasibility of this technique by acquiring 2D mechanical images of the neural plate *in situ* as the embryo was experiencing neurulation. We found that the neural plate tissue was continuously stiffened during NTC, which is consistent with previous observations of other species (Wiebe and Brodland, 2005; Zhou et al., 2009; Zhou et al., 2015; Barriga et al., 2018). The tissue stiffening is likely caused by the increased cell density and/or the accumulation of actomyosin contractility (Zhou et al., 2009; Galea et al., 2017; Barriga et al., 2018), which needs further investigation. Beyond that, we observed tissue stiffening is strongly correlated with the tissue thickening and bending.

Neurulation is a complex process involving cellular, molecular and biomechanical activities (Miller and Davidson, 2013; Nikolopoulou et al., 2017). While the genetic regulation and biochemical signaling have been extensively investigated, the biomechanical mechanism is less explored, and the underlying linkage between microscopic cellular/molecular activities and macroscopic morphogenesis is mostly unknown (Vijayraghavan and Davidson, 2017). Since our all-optical technique can directly quantify the tissue mechanics within intact live embryo, it can potentially open up new opportunities to better understand the role of biomechanical mechanism in the procedure of NTC. For example, a couple of crucial cellular activities including convergent extension (Wallingford et al., 2002), apical constriction (Sawyer et al., 2010), and interkinetic nuclear migration (Smith and Schoenwolf, 1987; Spear and Erickson, 2012) may together coordinate the observed tissue stiffening, thickening, and bending. The subcellular resolution of the Brillouin microscope will allow researchers to further investigate the role of these cellular behaviors in regulating tissue biomechanics. On the other hand, the mechanical cues can guide cell behaviors and cell fates through mechanotransduction during embryonic development (Miller and Davidson, 2013; Davis and Tapon, 2019); thus, the technique can also help understand the interaction between biochemical signaling and biomechanical cues. In addition, the closure of the neural tube is physically driven by both the generated force and the mechanical resistance of the tissue (Zhou et al., 2015; Galea et al., 2017; Moon and Xiong, 2021). The *in-situ* quantification of the mechanical properties can help decouple the roles of the force and the tissue mechanics thus allow better elucidation of the biomechanical interaction. Furthermore, the computational modeling is a powerful tool to understand the mechanism of morphogenesis (Davidson et al., 2010; Nishimura et al., 2012; Murisic et al., 2015). The time-lapse mechanical images of the neural plate tissue acquired by our technique can provide new input data for the simulation of neurulation.

Neural tube defects (NTDs) are among the commonest human birth diseases and regulated by both genetic and environmental factors (Blom et al., 2006; Copp et al., 2013). On the tissue level, NTDs arise from the physical failure of the neural tube due to the abnormal interaction of force generation and the mechanical properties of embryonic tissue. Recent work suggests that the NTD induced by gene mutation is associated with altered tissue biomechanics (Galea et al., 2017). Therefore, a quantitative tool for measuring tissue mechanics should allow researchers to attribute different NTDs to specific dysregulation of cellular mechanisms that cause the failure of the tissue closure, which could bridge the gap between genetic/environmental factors and tissue biomechanics and help the prevention of the diseases.

It is worth noting that, by definition, the high-frequency longitudinal modulus measured by Brillouin technique is different from the low-frequency or quasi-static Young’s modulus measured by conventional method such as AFM. However, for many biological materials, it is found that the two moduli change in the same direction in response to biological activities (Scarcelli and Yun, 2018; Wu et al., 2018). Therefore, with a careful calibration for specific material, one might interpret Brillouin data in terms of Young’s modulus. In this work, we used Brillouin shift to estimate the relative change of modulus by assuming the ratio of density and refractive index (*ρ/n*^2^) is constant. To directly quantify the Brillouin-derived modulus, Brillouin microscope can be combined with other technique that can measure the density and/or refractive index (Schlüßler et al., 2022). As the image depth increases, the strength of the Brillouin signal will drop depending on the transparency of the tissue. For chick embryo, the maximum penetration depth of our instrument is about 200 μm. Further improvement can be achieved by using a laser source with longer wavelength or wavefront correction technique based on adaptive optics (Edrei and Scarcelli, 2018).

## MATERIALS AND METHODS

### Eggs and *in ovo* culture

Fertilized white leghorn eggs were purchased from poultry farm of the University of Connecticut. For *in ovo* culturing, eggs were incubated at 37 °C under high humidity. Incubation hours follows Hamburger-Hamilton (HH) staging (i.e. 26-29 hrs of incubation to obtain HH-4 embryo) (Hamburger and Hamilton, 1992).

### Modified *ex ovo* culture of chick embryo

The *ex ovo* culture protocol was derived from Chapman *et al* and Schmitz *et al* (Chapman et al., 2001; Schmitz et al., 2016) with modifications for adapting to the Brillouin microscope. A 35 mm glass bottom dish with a 20 mm micro-well (Cellvis, D35-20-0-N) was used for *ex ovo* culture. A 1-inch metallic ring (Thorlabs, SM1RR) was covered with a single-layer Parafilm and a 1 cm ellipse hole was cut into the center and placed over the micro-well (inner bottom well of dish). To perform *ex ovo* culture, we used the filter paper carrier method to hold the blastoderm and vitelline membrane under tension to mimic the situation of in ovo culturing. The pre-cultured embryo around HH 4 was collected from the egg and then placed dorsal side down onto a culture dish filled with thin albumin harvested from the egg. Next, the dish was placed into an on-stage incubator (Warner Instruments, SA-20PLIXR-AL) for continuous culture. To ensure the ambient temperature of the embryo is about 37 °C, the heater (Warner Instruments, TC-344C) of the on-state incubator was set to 39 +/- 0.2 °C considering the heat dissipation from the underside of the stage which is open to the objective lens.

### Brillouin light scattering

Spontaneous Brillouin light scattering is the interactions between the incident light and inherent acoustic phonons inside a sample. The result of this interaction introduces a frequency shift (Brillouin shift) to the outgoing scattered light. The Brillouin shift *ω_B_* is defined as 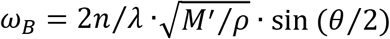, where *n* is refractive index of the material, *λ* is the laser wavelength, *M*′ is the longitudinal modulus that quantifies the mechanical property, *ρ* is the density, and *θ* is the collection angle of the scattered light. In our Brillouin microscope, backward scattered light was collected, yielding *θ* = 180 °.

### Estimation of Young’s modulus using measured Brillouin shift

For biological materials, an empirical relationship between Young’s modulus *E* and Brillouin-derived longitudinal modulus *M*′ has been established: log(*M*′) = *a* · log(*E*) + *b*, where the parameters *a* and *b* are material dependent (Scarcelli et al., 2015). Considering the relationship between *M*′ and Brillouin shift *ω_B_*, the relative change of Brillouin shift is related to that of Young’s modulus by Δ*E/E* =2/*a* · Δ*ω_B_/ω_B_*. For cells, the calibration against AFM shows *a* = 0.0671. Here, we used this calibrated relationship to estimate the relative change of Young’s modulus.

### Time-lapse Brillouin imaging

A confocal Brillouin microscope was used for all experiments. The detail of the instrumentation can be found in our recent report (Zhang and Scarcelli, 2021). Briefly, a single mode 660-nm continuous wave laser with power of ~30 mW was used as light source. The laser beam was focused into the sample by an objective lens (Olympus, 40×/0.6 NA) installed on an inverted microscope (Olympus, IX81), which yields a spot size of 0.7 μm × 0.7 μm × 2.6 μm. The backward scattered Brillouin signal was collected by the same objectives and analyzed by a two-stage VIPA (Light Machinery, 15 GHz FSR) based spectrometer, and the Brillouin spectrum was recorded by an EMCCD camera (Andor, iXon 897) with an exposure time of 0.05 s. Two dimensional Brillouin images were acquired by scanning the sample using a motorized stage (step size: 0.5 μm). The cross-section perpendicular to the anterior-posterior body axis was mapped by Brillouin microscope, and the averaged Brillouin shift of the neural plate region was used to represent the mechanical properties of the tissue. The Brillouin shift of the albumin is very close to that of the vitelline membrane, making it difficult to identify the boundary of neural plate tissue in the Brillouin image. To solve this issue, right before acquiring each Brillouin image, the embryo was temporarily transferred onto a different culture dish filled with Ringer’s solution (Thermo Scientific, BR0052G). As soon as the Brillouin measurement is done, the embryo was transferred back to the *ex ovo* culturing dish for continuous development. To acquire the bright-field images, a low magnification objective lens (Olympus, 4x/0.1) and a CMOS camera (Andor Neo) were used when the embryo was in the *ex ovo* culturing dish.

### Data acquisition and analysis

A home-built LabView (National Instruments, ver.2021) acquisition program was used to acquire both bright field images and the Brillouin spectra. For calibration of the spectrometer, Brillouin spectra of materials (water and methanol) with known Brillouin shifts were recorded and used to calculate the free spectral range and the pixel-to-frequency convention ratio. The Brillouin shift of each pixel was obtained by fitting the Brillouin spectrum to a Lorentzian function using MATLAB (MathWorks, R2021b). Two dimensional Brillouin image was reconstructed from the pixel vector. Sample size was chosen based on previous experience and to be reasonably large to demonstrate the feasibility of the technique.

## Acknowledgements

We thank Eric Frank for assisting the development of the LabView acquisition program.

## Data availability

All data supporting the findings of this study are available within the paper and its Supplementary Information files.

---

**Supplemental Figure 1.**
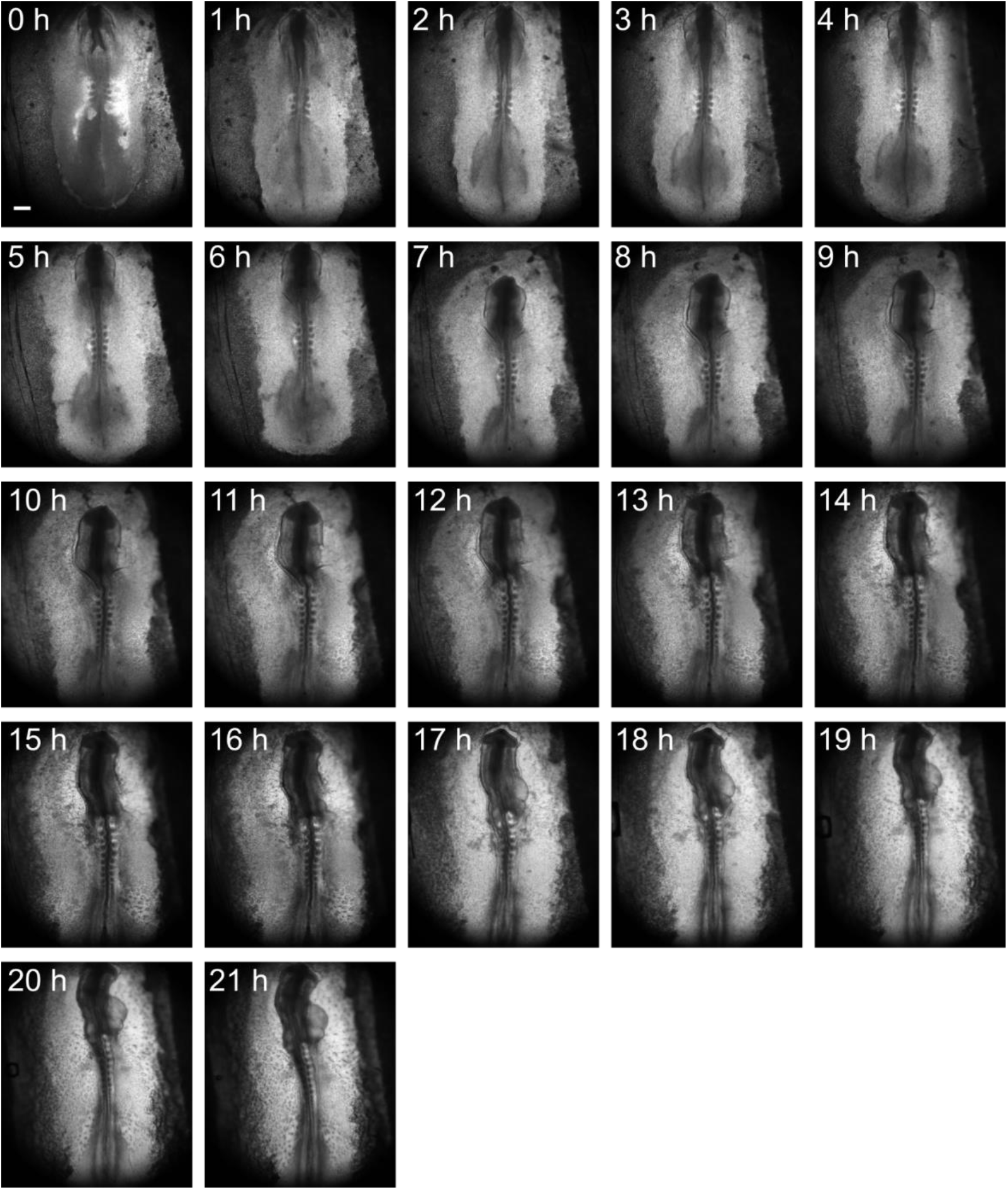
Brightfield time-lapse imaging of *ex ovo* cultured embryo without Brillouin measurement. Representative of continuous *ex ovo* development on thin albumin without laser illumination for 21 hrs. An HH8 embryo is extracted for *ex ovo* culture after 29 hrs of *in ovo* incubation (0 h). Embryo is identified as HH13+ at 21 h of *ex ovo* incubation (21 h) with a total incubation time of 50 hrs. Embryo develops according to the HH stages with identifiable markers at each stage. Each image taken 1 h apart. Scale bar is 300 μm.

**Supplemental Figure 2.**
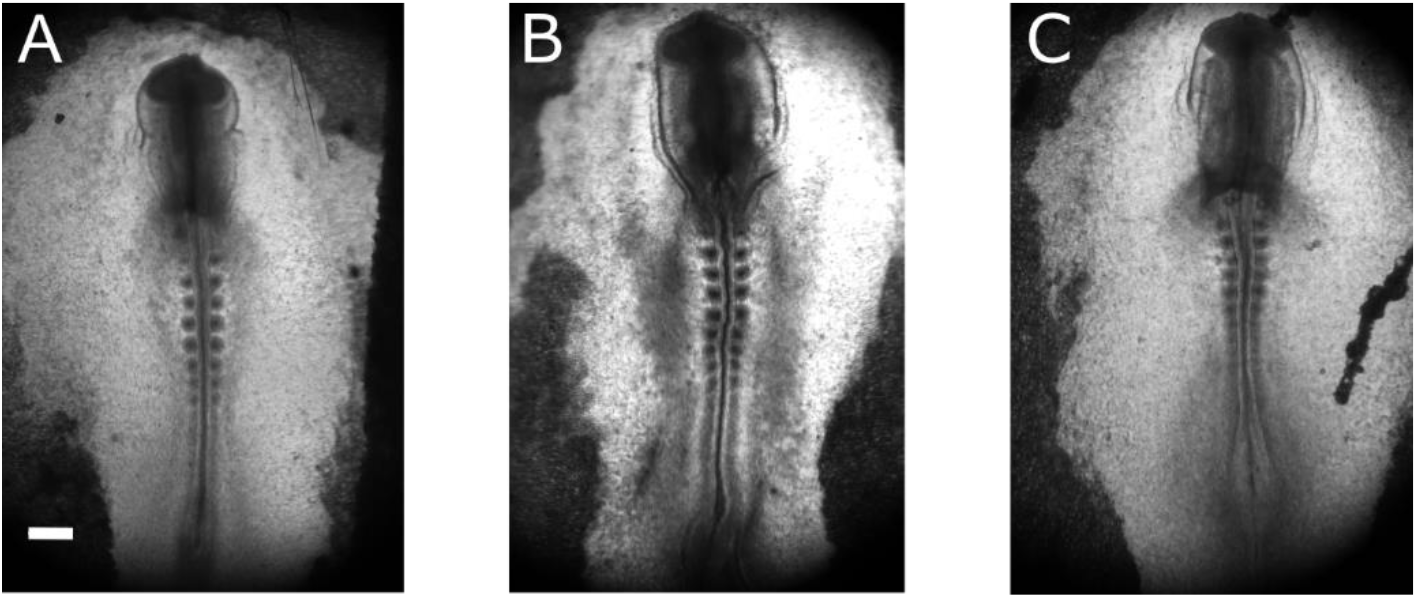
Comparison of all incubation and experimental methods reveal similar development. (A) HH 10 embryo after 39 hrs of continuous culture (30 hrs *in ovo* + 9 hrs *ex ovo*) on thin albumin without laser illumination. (B) HH 10 embryo after 38 hrs of time-lapse *ex ovo* culture (30 hrs *in ovo* + 8 hrs *ex ovo*) including transfer onto Ringer’s solution and exposure to laser illumination during Brillouin acquisition. (C) HH 10 embryo after 38 hrs of *in ovo* culture extracted for Brillouin mapping. All three embryos experienced similar development time following HH stages and display similar morphology with different methods of incubation and experimental setting. Scale bar is 300 μm.

**Supplemental Figure 3.**
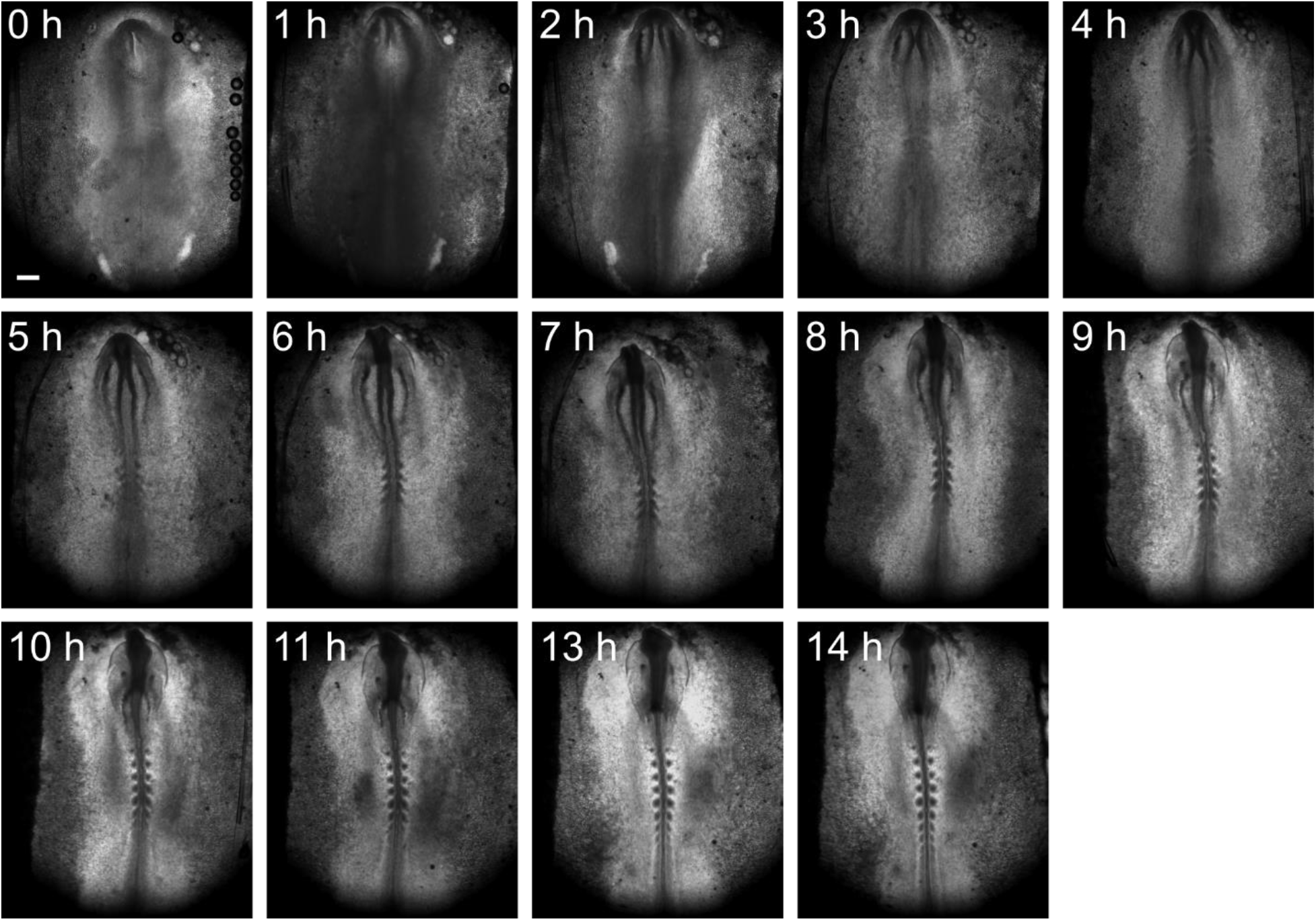
Brightfield time-lapse Brillouin imaging of *ex ovo* cultured embryo. Representative of 14 hrs of *ex ovo* development during Brillouin imaging, including transfer onto Ringer’s solution and exposure to laser illumination. An HH 8 embryo was extracted for *ex ovo* culture and time-lapse Brillouin imaging after 26 hrs *in ovo* incubation (0 h) and developed to an HH 10 embryo after 14 hrs of *ex ovo* incubation (14 h). Brillouin acquisition was conducted every 2 hrs with the initial acquisition at 0 h (HH8). This is to allow the embryo to develop on thin albumin between acquisition. Each image taken 1 h apart, 12 h not displayed. Scale bar is 300 μm.

## References

Barriga, Elias H, Franze, Kristian, Charras, Guillaume and Mayor, Roberto (2018) ‘Tissue stiffening coordinates morphogenesis by triggering collective cell migration in vivo’, Nature 554(7693): 523–527.

Beloussov, LV, Dorfman, JG and Cherdantzev, VG (1975) ‘Mechanical stresses and morphological patterns in amphibian embryos’, Development 34(3): 559–574.

Blom, Henk J, Shaw, Gary M, den Heijer, Martin and Finnell, Richard H (2006) ‘Neural tube defects and folate: case far from closed’, Nature Reviews Neuroscience 7(9): 724.

Boyd, Robert W (2003) Nonlinear optics: Academic press.

Campas, O. (2016) ‘A toolbox to explore the mechanics of living embryonic tissues’, Seminars in Cell and Developmental Biology 55: 119–130.

Campàs, Otger, Mammoto, Tadanori, Hasso, Sean, Sperling, Ralph A, O’connell, Daniel, Bischof, Ashley G, Maas, Richard, Weitz, David A, Mahadevan, Lakshminarayanan and Ingber, Donald E (2014) ‘Quantifying cell-generated mechanical forces within living embryonic tissues’, Nature Methods 11(2): 183–189.

Chapman, Susan C, Collignon, Jérôme, Schoenwolf, Gary C and Lumsden, Andrew (2001) ‘Improved method for chick whole-embryo culture using a filter paper carrier’, Developmental dynamics 220(3): 284–289.

Chevalier, Nicolas R, Gazguez, Elodie, Dufour, Sylvie and Fleury, Vincent (2016) ‘Measuring the micromechanical properties of embryonic tissues’, Methods 94: 120–128.

Colas, Jean-François and Schoenwolf, Gary C (2001) ‘Towards a cellular and molecular understanding of neurulation’, Developmental dynamics: an official publication of the American Association of Anatomists 221(2): 117–145.

Copp, Andrew J, Greene, Nicholas DE and Murdoch, Jennifer N (2003) ‘The genetic basis of mammalian neurulation’, Nature Reviews Genetics 4(10): 784.

Copp, Andrew J, Stanier, Philip and Greene, Nicholas DE (2013) ‘Neural tube defects: recent advances, unsolved questions, and controversies’, The Lancet Neurology 12(8): 799–810.

Davidson, Lance A, Joshi, Sagar D, Kim, Hye Young, Von Dassow, Michelangelo, Zhang, Lin and Zhou, Jian (2010) ‘Emergent morphogenesis: elastic mechanics of a self-deforming tissue’, Journal of Biomechanics 43(1): 63–70.

Davis, John Robert and Tapon, Nicolas (2019) ‘Hippo signalling during development’, Development 146(18): dev167106.

Edrei, Eitan and Scarcelli, Giuliano (2018) ‘Brillouin micro-spectroscopy through aberrations via sensorless adaptive optics’, Applied physics letters 112(16): 163701.

Franze, Kristian (2011) ‘Atomic force microscopy and its contribution to understanding the development of the nervous system’, Current Opinion in Genetics and Development 21(5): 530–537.

Galea, G. L., Cho, Y. J., Galea, G., Mole, M. A., Rolo, A., Savery, D., Moulding, D., Culshaw, L. H., Nikolopoulou, E., Greene, N. D. E. et al. (2017) ‘Biomechanical coupling facilitates spinal neural tube closure in mouse embryos’, Proceedings of the National Academy of Sciences of the United States of America 114(26): E5177–e5186.

Hamburger, Viktor and Hamilton, Howard L (1992) ‘A series of normal stages in the development of the chick embryo’, Developmental Dynamics 195(4): 231–272.

Heer, Natalie C and Martin, Adam C (2017) ‘Tension, contraction and tissue morphogenesis’, Development 144(23): 4249–4260.

Koehl, MAR (1990) Biomechanical approaches to morphogenesis Seminars in Developmental Biology, vol. 1.

Lowery, Laura Anne and Sive, Hazel (2004) ‘Strategies of vertebrate neurulation and a re-evaluation of teleost neural tube formation’, Mechanisms of Development 121(10): 1189–1197.

Marrese, Marica, Antonovaite, Nelda, Nelemans, Ben KA, Smit, Theodoor H and Iannuzzi, Davide (2019) ‘Micro-indentation and optical coherence tomography for the mechanical characterization of embryos: Experimental setup and measurements on chicken embryos’, Acta Biomaterialia 97: 524–534.

Miller, Callie Johnson and Davidson, Lance A (2013) ‘The interplay between cell signalling and mechanics in developmental processes’, Nature Reviews Genetics 14(10): 733–744.

Moon, Lauren D and Xiong, Fengzhu (2021) Mechanics of neural tube morphogenesis Seminars in Cell and Developmental Biology: Elsevier.

Murisic, Nebojsa, Hakim, Vincent, Kevrekidis, Ioannis G, Shvartsman, Stanislav Y and Audoly, Basile (2015) ‘From discrete to continuum models of three-dimensional deformations in epithelial sheets’, Biophysical Journal 109(1): 154–163.

Nikolopoulou, Evanthia, Galea, Gabriel L, Rolo, Ana, Greene, Nicholas DE and Copp, Andrew J (2017) ‘Neural tube closure: cellular, molecular and biomechanical mechanisms’, Development 144(4): 552–566.

Nishimura, Tamako, Honda, Hisao and Takeichi, Masatoshi (2012) ‘Planar cell polarity links axes of spatial dynamics in neural-tube closure’, Cell 149(5): 1084–1097.

Prevedel, Robert, Diz-Muñoz, Alba, Ruocco, Giancarlo and Antonacci, Giuseppe (2019) ‘Brillouin microscopy: an emerging tool for mechanobiology’, Nature Methods 16(10): 969–977.

Raghunathan, Raksha, Zhang, Jitao, Wu, Chen, Rippy, Justin, Singh, Manmohan, Larin, Kirill V and Scarcelli, Giuliano (2017) ‘Evaluating biomechanical properties of murine embryos using Brillouin microscopy and optical coherence tomography’, Journal of Biomedical Optics 22(8): 086013.

Savin, Thierry, Kurpios, Natasza A, Shyer, Amy E, Florescu, Patricia, Liang, Haiyi, Mahadevan, L and Tabin, Clifford J (2011) ‘On the growth and form of the gut’, Nature 476(7358): 57–62.

Sawyer, Jacob M, Harrell, Jessica R, Shemer, Gidi, Sullivan-Brown, Jessica, Roh-Johnson, Minna and Goldstein, Bob (2010) ‘Apical constriction: a cell shape change that can drive morphogenesis’, Developmental Biology 341(1): 5–19.

Scarcelli, Giuliano, Kling, Sabine, Quijano, Elena, Pineda, Roberto, Marcos, Susana and Yun, Seok Hyun (2013) ‘Brillouin microscopy of collagen crosslinking: noncontact depth-dependent analysis of corneal elastic modulus’, Investigative Ophthalmology and Visual Science 54(2): 1418–1425.

Scarcelli, Giuliano, Polacheck, William J., Nia, Hadi T., Patel, Kripa, Grodzinsky, Alan J., Kamm, Roger D. and Yun, Seok Hyun (2015) ‘Noncontact three-dimensional mapping of intracellular hydro-mechanical properties by Brillouin microscopy’, Nature Methods 12(12): 1132–1134.

Scarcelli, Giuliano and Yun, Seok Hyun (2018) ‘Reply to ‘Water content, not stiffness, dominates Brillouin spectroscopy measurements in hydrated materials’’, Nature Methods 15(8): 562.

Schlüßler, Raimund, Kim, Kyoohyun, Nötzel, Martin, Taubenberger, Anna, Abuhattum, Shada, Beck, Timon, Müller, Paul, Maharana, Shovamaye, Cojoc, Gheorghe and Girardo, Salvatore (2022) ‘Correlative all-optical quantification of mass density and mechanics of sub-cellular compartments with fluorescence specificity’, Elife 11: e68490.

Schmitz, Manuel, Nelemans, Ben KA and Smit, Theodoor H (2016) ‘A submerged filter paper sandwich for long-term ex ovo time-lapse imaging of early chick embryos’, JoVE (Journal of Visualized Experiments)(118): e54636.

Schoenwolf, Gary C and Smith, Jodi L (1990) ‘Mechanisms of neurulation: traditional viewpoint and recent advances’, Development 109(2): 243–270.

Smith, Jodi L and Schoenwolf, Gary C (1987) ‘Cell cycle and neuroepithelial cell shape during bending of the chick neural plate’, The Anatomical Record 218(2): 196–206.

Spear, Philip C and Erickson, Carol A (2012) ‘Interkinetic nuclear migration: a mysterious process in search of a function’, Development, growth & differentiation 54(3): 306–316.

Vijayraghavan, Deepthi S and Davidson, Lance A (2017) ‘Mechanics of neurulation: From classical to current perspectives on the physical mechanics that shape, fold, and form the neural tube’, Birth defects research 109(2): 153–168.

Wallingford, John B, Fraser, Scott E and Harland, Richard M (2002) ‘Convergent extension: the molecular control of polarized cell movement during embryonic development’, Developmental Cell 2(6): 695–706.

Wallingford, John B, Niswander, Lee A, Shaw, Gary M and Finnell, Richard H (2013) ‘The continuing challenge of understanding, preventing, and treating neural tube defects’, Science 339(6123): 1222002.

Welte, Michael A, Gross, Steven P, Postner, Marya, Block, Steven M and Wieschaus, Eric F (1998) ‘Developmental regulation of vesicle transport in Drosophila embryos: forces and kinetics’, Cell 92(4): 547–557.

Wen, Jun, Liu, Jun, Lau, Kimberly, Liu, Haijiao, Hopyan, Sevan and Sun, Yu (2015) Automated micro-aspiration of mouse embryo limb bud tissue 2015 IEEE International Conference on Robotics and Automation (ICRA): IEEE.

Wiebe, Colin and Brodland, G Wayne (2005) ‘Tensile properties of embryonic epithelia measured using a novel instrument’, Journal of Biomechanics 38(10): 2087–2094.

Wilde, Jonathan J, Petersen, Juliette R and Niswander, Lee (2014) ‘Genetic, epigenetic, and environmental contributions to neural tube closure’, Annual Review of Genetics 48: 583–611.

Wisniewski, Emily O, Mistriotis, Panagiotis, Bera, Kaustav, Law, Robert A, Zhang, Jitao, Nikolic, Milos, Weiger, Michael, Parlani, Maria, Tuntithavornwat, Soontorn and Afthinos, Alexandros (2020) ‘Dorsoventral polarity directs cell responses to migration track geometries’, Science Advances 6(31): eaba6505.

Wu, Pei-Jung, Kabakova, Irina V, Ruberti, Jeffrey W, Sherwood, Joseph M, Dunlop, Iain E, Paterson, Carl, Török, Peter and Overby, Darryl R (2018) ‘Water content, not stiffness, dominates Brillouin spectroscopy measurements in hydrated materials’, Nature Methods 15(8): 561.

Zhang, Jitao, Alisafaei, Farid, Nikolić, Miloš, Nou, Xuefei A, Kim, Hanyoup, Shenoy, Vivek B and Scarcelli, Giuliano (2020) ‘Nuclear mechanics within intact cells is regulated by cytoskeletal network and internal nanostructures’, Small 16(18): 1907688.

Zhang, Jitao, Raghunathan, Raksha, Rippy, Justin, Wu, Chen, Finnell, Richard H, Larin, Kirill V and Scarcelli, Giuliano (2018) ‘Tissue biomechanics during cranial neural tube closure measured by Brillouin microscopy and optical coherence tomography’, Birth defects research 111: 991–998.

Zhang, Jitao and Scarcelli, Giuliano (2021) ‘Mapping mechanical properties of biological materials via an add-on Brillouin module to confocal microscopes’, Nature Protocols 16(2): 1251–1275.

Zhou, J., Kim, H. Y. and Davidson, L. A. (2009) ‘Actomyosin stiffens the vertebrate embryo during crucial stages of elongation and neural tube closure’, Development 136(4): 677–688.

Zhou, Jian, Pal, Siladitya, Maiti, Spandan and Davidson, Lance A (2015) ‘Force production and mechanical accommodation during convergent extension’, Development 142(4): 692–701.

